# TraceTrack, an Open-Source Software for Batch Processing, Alignment and Visualization of Sanger Sequencing Chromatograms

**DOI:** 10.1101/2022.07.28.501824

**Authors:** Kveta Brazdilova, David Prihoda, Quynh Ton, Heath Klock, Danny A. Bitton

**Affiliations:** R&D Informatics Solutions, MSD Czech Republic s.r.o., Prague, Czech Republic; Department of Informatics and Chemistry, Faculty of Chemical Technology, University of Chemistry and Technology, Prague, Czech Republic; Protein Sciences, MRL, Merck & Co., Inc., Boston, Massachusetts, USA

## Abstract

**Background:** Despite the advent of Next Generation Sequencing technology and its widespread applications, Sanger sequencing remains instrumental for molecular biology subcloning work in biological and medical research and indispensable for drug discovery campaigns. Although Sanger sequencing technology has been long established, existing software for processing and visualization of trace file chromatograms are limited in terms of functionality, scalability, and availability for commercial use.

**Results:** To fill this gap, we developed TraceTrack, an open-source web application tool for batch alignment, analysis and visualization of Sanger trace files. TraceTrack offers high throughput matching of trace files to reference sequences, rapid identification of mutations and an intuitive chromatogram analysis. Comparative analysis between TraceTrack and existing software tools highlights the advantages of TraceTrack with regards to batch processing, visualization and export functionalities. TraceTrack is available at https://github.com/Merck/TraceTrack and also at https://tracetrack.dichlab.org as a web application.

**Conclusion:** TraceTrack is a web application for batch processing and visualization of Sanger trace file chromatograms that meets the increasing demand of industrial sequence validation workflows in pharmaceutical settings.

## Background

Although Next Generation Sequencing (NGS) technologies have revolutionized biomedical research and high-throughput methodologies in genomics and transcriptomics (1, 2), Sanger sequencing still plays a pivotal role in sequence validation primarily due to its simplicity, accuracy, and cost-effectiveness (3). Even where NGS is widely used, Sanger sequencing remains important for validation of variants, completing hard to sequence regions (4, 5), identifying pathogens (6, 7) as well as for Short Tandem Repeat (STR) analysis (8). In pharmaceutical research, Sanger sequencing is routinely and extensively used for primer walking (2, 9), cloning junction verifications and point mutation detection, supporting a wide-range of high-throughput pipelines for protein design, biocatalysis and antibody discovery, to name just a few. Manual interpretation of Sanger sequencing chromatograms and their comparison to reference sequences may take an experienced researcher approximately 5 minutes to complete for a single sample. However, when processing multiple batches of sequencing data in 96-well plates, chromatogram matching and analysis becomes a tedious and error-prone process. Thus, the heavy workload of sequence analysis and validation in drug discovery workflows demands the development of automated, high-throughput Sanger-based validation tools that can dramatically reduce manual curation of sequences, and consequently save cost and time.

In this regard, many applications for automated matching and alignment of Sanger trace files to reference sequences have been reported to date (10-17). Nevertheless, many of the published software tools are no longer being supported (10-12), not freely available for widespread commercial use (13-15) or cannot handle batch processing, therefore, are unable to support large-scale chromatogram analysis in pharmaceutical settings. A notable example is Tracy (16), a recent open-source solution that represents a step forward for automating chromatogram analysis. Tracy features advanced functionalities for genome assembly, base-calling and trace file alignment through an intuitive user interface, yet it remains unsuitable for high throughput analysis, since trace files can only be processed in bulk using a command-line interface, which limits is use by non-tech-savvy users. Similarly, the recently published ‘R’ package “sangeranalyseR” (17) offers an advanced toolbox to streamline and speed up chromatogram analysis, nevertheless it requires significant knowledge of the R programming language.

To address this unmet need we developed TraceTrack, an open-source web application for batch alignment of trace files, mutation detection and chromatogram visualization that can significantly reduce the time of sequence validation (Figure 1). Via an intuitive user-interface TraceTrack enables simultaneous matching and alignment of multiple trace files to multiple reference sequences. Each resultant alignment is displayed separately with highlighted sequence variations. Original chromatograms can also be interrogated directly. TraceTrack is an extensible software tool not only enabling sequence validation at speed and scale, but also aiming to encourage scientists to add additional features that can further reduce manual work and consequently accelerate drug discovery campaigns.

**Figure 1.**
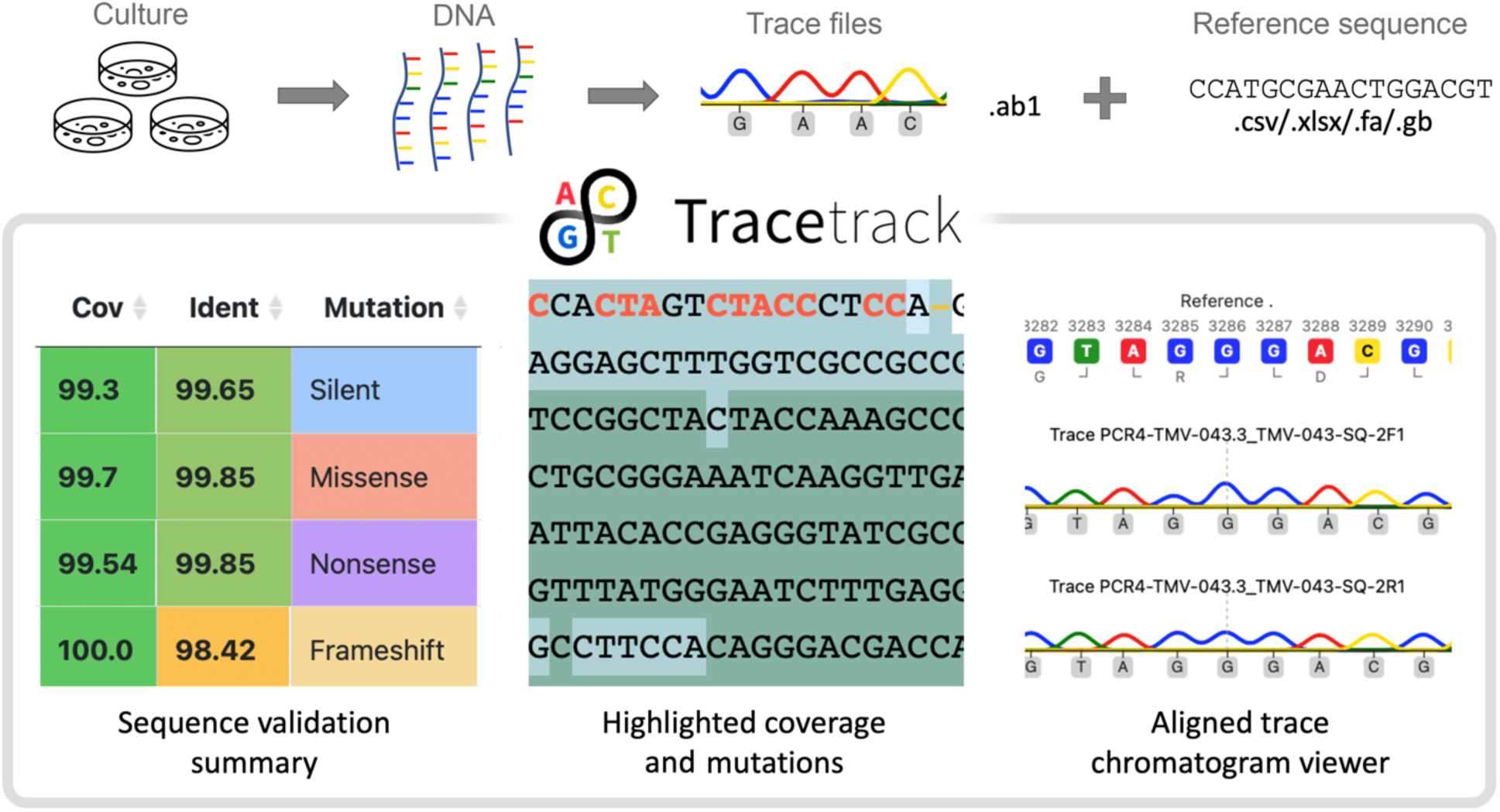
TraceTrack enables high throughput matching of Sanger chromatogram files and reference sequences. (Top) TraceTrack is routinely and extensively used to validate sequences in subcloning work. Sanger sequencing trace files (.ab1 file format) are matched with reference sequences (.csv, .xslx, .fa, .gb file formats) via systematic multiple sequence alignments. (Bottom) For each resultant alignment TraceTrack reports consensus sequence alongside percentage coverage, identity and the type of detected mutations as well as enables users to interrogate trace files via a dedicated trace viewer.

## Implementation

TraceTrack features a user-friendly interface and an extensible computational backend which are described in detail in the following sections.

### Application Overview

The web application is composed of three main components: (i) an html and JavaScript user interface with a Flask backend, (ii) a worker with a Celery asynchronous task queue, where computation takes place, (iii) a Redis in-memory database that is used to temporarily store information about the task and its respective results. Each of these components can be run separately in a terminal, or all three parts can be run together in a single Docker container. TraceTrack offers an easy and versatile deployment of all components on a local machine or on a remote server. Alternatively, deployment can be further simplified by running the application with synchronous tasks, thereby excluding the task queue and the database components.

TraceTrack employs BioPython (18) built-in functions for sequence manipulations and for extracting information from ab1 trace files. Yet, it offers numerous new custom classes and functions for storing trace and reference sequences as well as for sequence alignment. To perform multiple sequence alignment (MSA) of traces and references sequences, TraceTrack utilises the Clustal Omega algorithm (19) via the Bio.Align package (18). In cases where only two sequences are being aligned the same algorithm is used for consistency. The tool architecture and basic workflows are illustrated in Figure 2.

**Figure 2.**
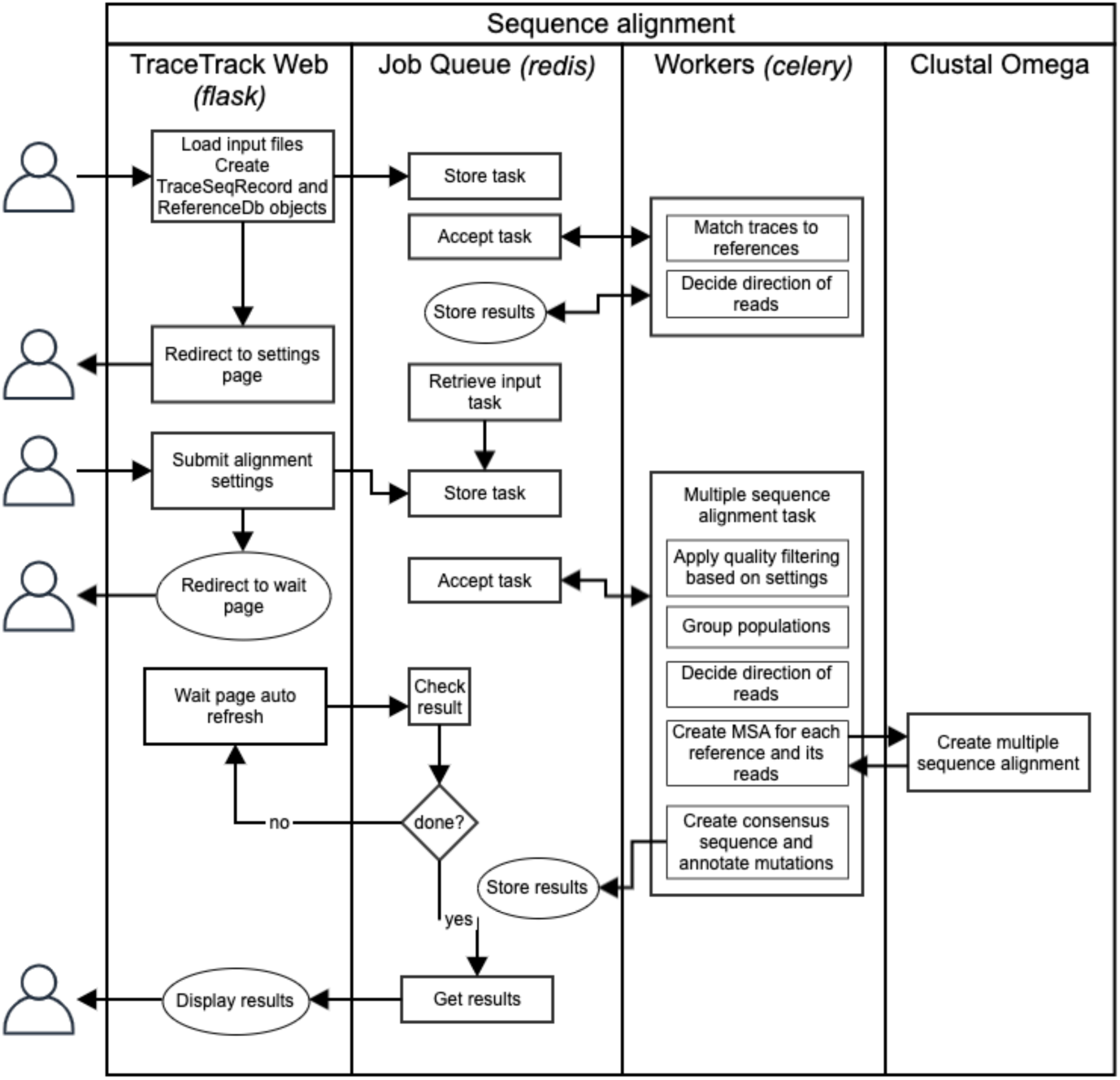
Tool architecture. In the basic workflow, the user uploads a set of trace files and a sheet containing reference sequences, which are queued as a task. Next, the user inputs settings, which are stored as another task. This task is accepted from the queue by the worker and corresponding objects are created from trace and reference sequences. These are then quality trimmed, grouped and reversed where necessary, before the resulting multiple sequence alignment is produced for each reference sequence with its respective traces. The result is then stored. The frontend checks for the result regularly and when it is available, displays it to the user.

### File input

TraceTrack’s input page displays buttons for uploading trace and reference sequence files (Figure 3A). Trace files can be uploaded in ab1 format or as a zipped archive with multiple ab1 files. TraceTrack accepts a single file with reference sequences in one of the following formats: .xlsx, .csv, .fasta or .gb (GenBank). Hundreds of traces and reference sequences can be uploaded and processed simultaneously, and users can optionally define distinct sub-groups of traces to be aligned separately.

**Figure 3.**
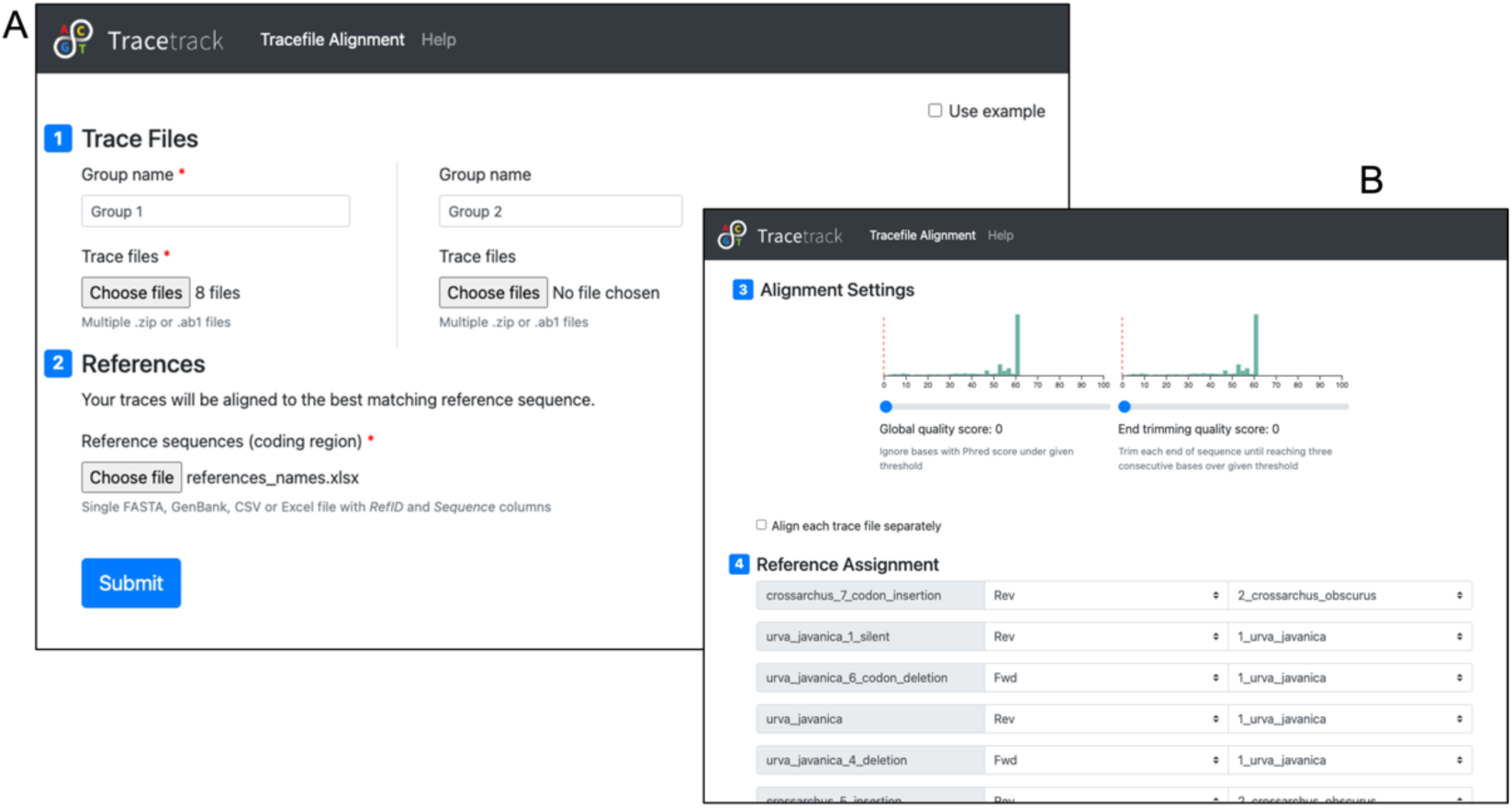
TraceTrack input page (A) Users can upload hundreds of Sanger trace files and reference sequences for matching. (B) Settings page with options for choosing references and read directions. The data analysed in these screenshots are sequencing reads from *Urva javanica* and *Crossarchus obscurus* obtained from the Barcode of Life Database (20)

### File processing

Once user uploads the trace and reference sequence files, TraceTrack generates a sequence list and a reference database in preparation for the subsequent matching and MSAs. When the reference is provided in GenBank format, TraceTrack simply extracts the coding sequence (CDS) features under default settings. In any other case, the entire sequence is considered as the coding sequence, and translation begins at the first codon in the sequence and ends with the last triplet of the sequence.

The software inspects the trace chromatograms from each file for mixed peaks. Each chromatogram is composed of four traces that correspond to each of the DNA nucleotides (A, T, G and C). In principle only a single peak in one of the traces at a given position should be identified. In cases where more than one peak is present, TraceTrack detects it, but it reports it only when the peaks are detected in high signal-to-noise regions (based on a data-derived threshold) as opposed to mixed peaks in low-complexity regions that are ignored throughout. To define a mixed peak, the area of the secondary peak must be at least ‘*f*’ times the area of the main peak and the secondary peak must be at least ‘*f*’ times the height of the main peak, where ‘*f*’ is a number between 0 and 1 (as derived from sample data; default 0.15) and the secondary trace is concave around the centre of the main peak.

Finally, TraceTrack pre-assigns all trace files to their respective reference sequence, either by sequence similarity or by matching filenames.

### Reference Assignment and Alignment Settings

TraceTrack settings page displays a list of trace files and their pre-assigned reference sequences as well as several settings options (Figure 3B). Each uploaded trace file is assigned to the best matching reference sequence, as follows. If the reference ID is contained in the trace file name, the trace file is assigned to it. Otherwise, the trace file is aligned to all references in both directions and the match with the highest score is chosen. Sequencing direction is determined automatically by evaluating read matches in both directions. These automatic reference assignments can be refined by the user using drop-down lists on the settings page. Trace sequences may also be filtered or trimmed according to user-defined quality or trimming thresholds. When a base quality threshold is set, the software discards all positions with a lower quality. When an end trimming threshold is set, the tool trims the ends of each trace sequence until three consecutive bases with quality higher than the threshold are encountered. This is intended to remove ends of reads with low quality, even when they contain some bases passing the quality threshold. TraceTrack ignores ambiguous base calls (“N”s) throughout.

### Multiple Sequence Alignment, Consensus Sequence and Mutation Calls

As mentioned earlier, TraceTrack employs Clustal Omega to perform MSA. For each reference within a given group the MSA is created separately with all its corresponding trace files. The resultant aligned sequences are then used to generate a consensus sequence, according to the following principles: (i) TraceTrack calls point mutations, insertions and deletions with respect to the reference sequence only if all reads agree, (ii) in case only some traces contain an insertion, an ambiguous insertion character is displayed (“?”) and the base is not considered as a viable sequence position, (iii) in all other cases when the reads do not agree, the reference base is kept and the number of ignored reads is shown as reduced read coverage.

Once consensus sequence is defined, TraceTrack translates the CDS and classifies mutations as silent (same amino acid), missense (different amino acid), nonsense (produces a stop codon) or frameshift (caused by insertions and deletions).

Since TraceTrack translates the coding sequence continuously from the beginning, an insertion or deletion of a number of bases not divisible by three leads to a frameshift. All the shifted positions are displayed in a different colour and the translation continues in the new frame.

### Alignment and Results

After all trace files are assigned to their corresponding reference sequences, TraceTrack performs the alignments, generates consensus sequences, calls the mutations, and displays the results in a sortable table containing one alignment per row (Figure 4A). The rows are labelled with a reference ID and contain the following information: percentage of sequence coverage and percentage identity, numbers of different mutation types in the consensus sequence, and the number and names of aligned trace files.

**Figure 4.**
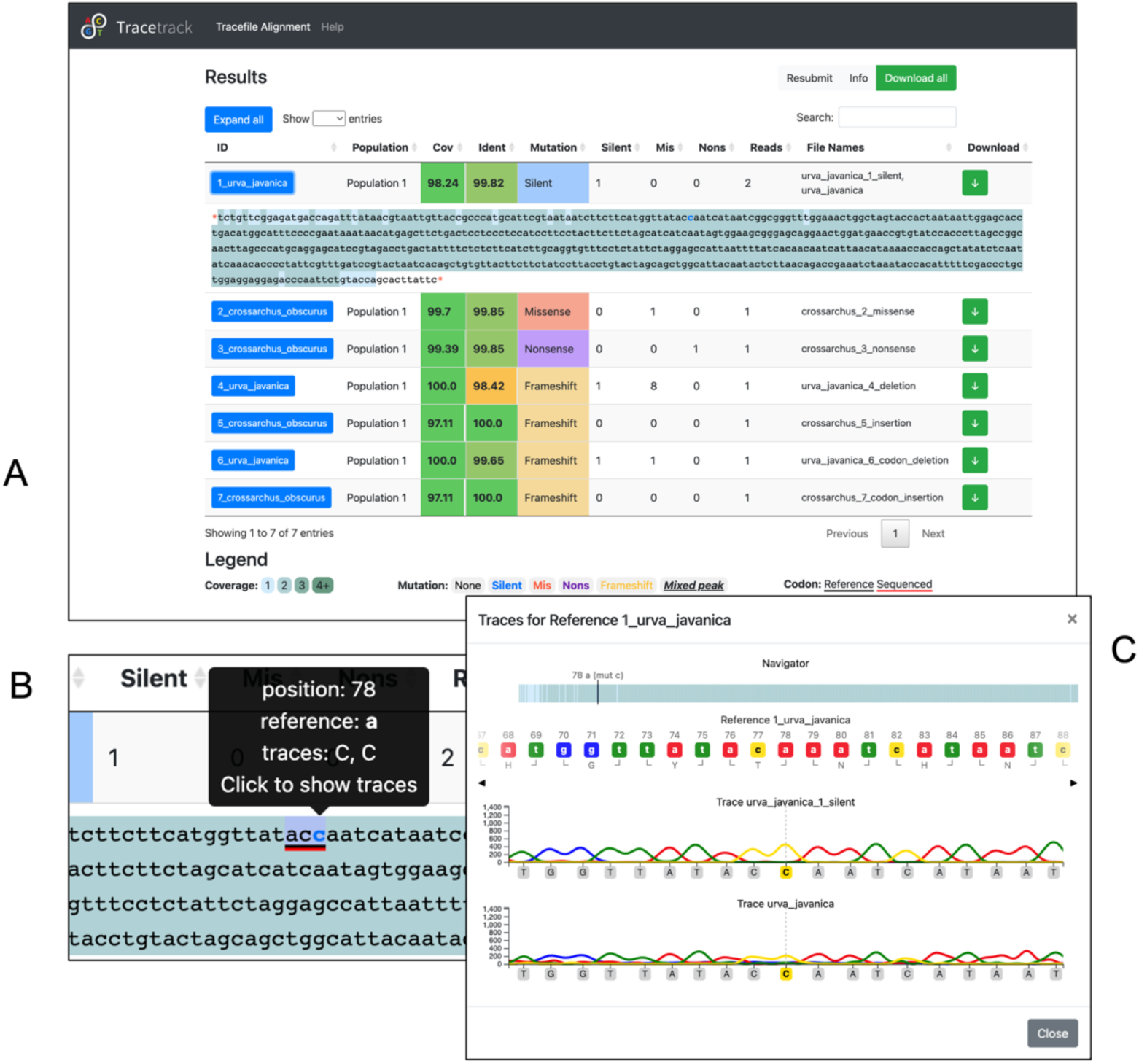
TraceTrack results page (A) Table of trace and reference sequence alignments (B) Alignment detail with colour-coded coverage and mutations (C) Trace viewer showing the reference sequence and traces aligned to it.

Each row can be expanded to show the consensus sequence with colour-coded coverage and respective mutations (Figure 4B). Background colour reflects coverage, text colour represents mutation type. Additional information about each position is provided by the tooltips widget, including the position and the base for each aligned trace sequence at that position.

### Chromatogram Visualization

When the user clicks any position in the alignment, a trace viewer appears. The reference sequence is displayed at the top and all trace chromatograms and sequences are shown underneath (Figure 4C). Amino acid translations are shown for the reference sequence and differences relative to the consensus sequence are highlighted. An interactive navigation bar can be used to navigate to specific mutations. The viewer can also be navigated using arrow buttons or by clicking any position in the reference or trace sequences.

### Exporting Results

A summary report can be downloaded as an interactive spreadsheet. The first sheet contains the overview table with links to subsequent sheets, which contain individual alignments. Both the reference and consensus sequences are displayed, as well as each trace sequence, along with the corresponding amino acid translations. A second sheet is also provided for each alignment with a list of all mismatching positions and regions with zero coverage. The sequences can be easily navigated by clicking mutation positions. If desired, each alignment can also be downloaded separately using a download button in the corresponding table row.

## Results

Taken together, TraceTrack enables batch processing of trace files and their alignment to multiple reference sequences as well as streamlines the inspection of chromatograms via a user-friendly web application. A comparative analysis to existing commercial or freely available tools highlights TraceTrack’s advantages and limitations (Table 1). The tool is flexible, it can run both on a web server as well as locally on a personal computer, and for quick access, TraceTrack is also available as a web application at https://tracetrack.dichlab.org. The main advantage of TraceTrack compared to other available open-source tools is the ability to process large numbers of both trace files and reference sequences simultaneously. In terms of computational performance, matching and aligning of a hundred of trace files to their respective reference sequences takes about a minute on a standard laptop computer. All the while the user is guided by a convenient graphical interface with no need for command line use or any knowledge of scripting.

**Table 1.**
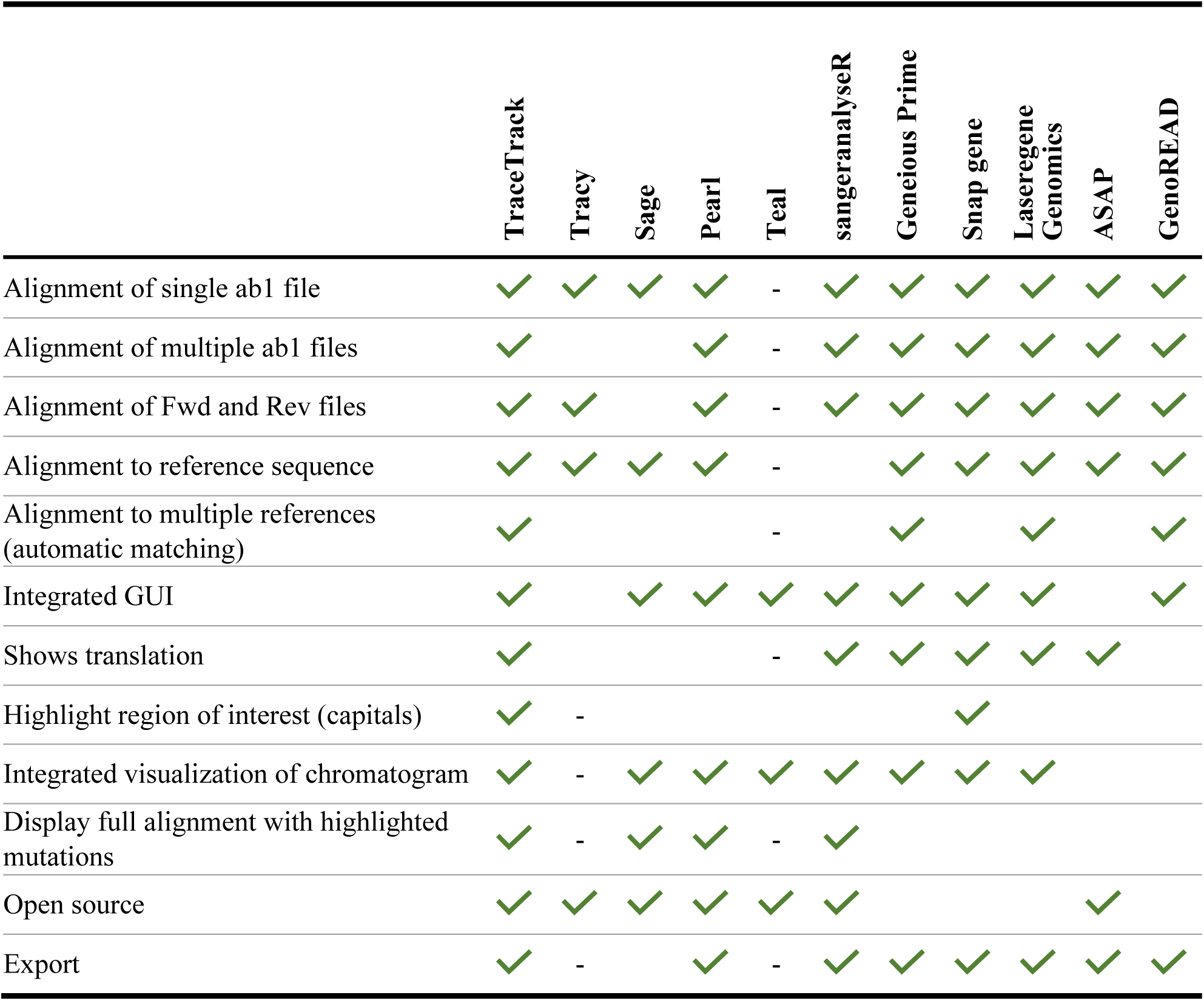
Comparison between TraceTrack and freely available and commercial tools

|  | TraceTrack | Tracy | Sage | Pearl | Teal | sangeranalyseR | Geneious Prime | Snap gene | Laseregene Genomics | ASAP | GenoREAD |
| --- | --- | --- | --- | --- | --- | --- | --- | --- | --- | --- | --- |
| Alignment of single ab1 file | ✓ | ✓ | ✓ | ✓ | - | ✓ | ✓ | ✓ | ✓ | ✓ | ✓ |
| Alignment of multiple ab1 files | ✓ |  |  | ✓ | - | ✓ | ✓ | ✓ | ✓ | ✓ | ✓ |
| Alignment of Fwd and Rev files | ✓ | ✓ |  | ✓ | - | ✓ | ✓ | ✓ | ✓ | ✓ | ✓ |
| Alignment to reference sequence | ✓ | ✓ | ✓ | ✓ | - |  | ✓ | ✓ | ✓ | ✓ | ✓ |
| Alignment to multiple references (automatic matching) | ✓ |  |  |  | - |  | ✓ |  | ✓ |  | ✓ |
| Integrated GUI | ✓ |  | ✓ | ✓ | ✓ | ✓ | ✓ | ✓ | ✓ |  | ✓ |
| Shows translation | ✓ |  |  |  | - | ✓ | ✓ | ✓ | ✓ | ✓ |  |
| Highlight region of interest (capitals) | ✓ | - |  |  |  |  |  | ✓ |  |  |  |
| Integrated visualization of chromatogram | ✓ | - | ✓ | ✓ | ✓ | ✓ | ✓ | ✓ | ✓ |  |  |
| Display full alignment with highlighted mutations | ✓ | - | ✓ | ✓ | - | ✓ |  |  |  |  |  |
| Open source | ✓ | ✓ | ✓ | ✓ | ✓ | ✓ |  |  |  | ✓ |  |
| Export | ✓ | - |  | ✓ | - | ✓ | ✓ | ✓ | ✓ | ✓ | ✓ |

Chromatograms annotated by the sequence and translation can be directly viewed within the application, providing an easy way to compare trace files to each other, or to the reference, and viewing changes at the amino acid level. A report can be downloaded for the entire analysis or for separate alignments.

Despite the advantages listed above it is important to note TraceTrack’s limitations. The use of an out-of-the-box multiple sequence alignment algorithm might be rigid and less flexible when it comes to handling indels and other edge cases in the alignment, providing an opportunity to further improve read alignment using more advanced read mapping algorithms. TraceTrack could further be extended by adding functionality to adjust trimming of trace sequences manually and edit base calls. Currently, trace sequences can only be trimmed based on quality thresholds and the sequence is immutable.

## Conclusion

As Sanger sequencing is still a widely used for biological and biomedical research, there is a need to find automated solutions to common tasks. The available open-source tools for aligning and inspecting trace files were scarce and lacked functionality for batch processing. Therefore, we developed TraceTrack, an application to bridge this gap and facilitate many areas of research, including protein design, biocatalysis and therapeutic antibody discovery. Since the tool is fully open-source, there is potential for updates and extensions not only by the authors, but also the greater scientific community.

## List of Abbreviations

MSA: Multiple sequence alignment
NGS: Next generation sequencing
STR: Short tandem repeat

## Availability of data and materials

All code for this publication is available in the following GitHub repository: https://github.com/Merck/TraceTrack.

## Author Contributions

DB and HK conceived the study. KB and DP developed and implemented the entire application. DP and DB supervised the study. QT advised and tested the tool throughout its development. DB and KB wrote the manuscript. All authors contributed to the final draft. All authors reviewed and approved the final draft.

## Acknowledgments

We thank Jens Christensen, Vincent Antonucci, and Carol A. Rohl for supporting this work. We are immensely grateful to Arthur Fridman, Charles Tilford, and Nick Mukhitov for their advice and stimulating discussions. We also thank Anja Muzdalo for reviewing the manuscript and Martin Spale for preparing the tool for open-source.

